# The distribution of deleterious genetic variation in human populations

**DOI:** 10.1101/005330

**Authors:** Kirk E. Lohmueller

**Affiliations:** Department of Ecology and Evolutionary Biology, Interdepartmental Program in Bioinformatics, University of California, Los Angeles, CA, 90095

## Abstract

Population genetic studies suggest that most amino-acid changing mutations are deleterious. Such mutations are of tremendous interest in human population genetics as they are important for the evolutionary process and may contribute risk to common disease. Genomic studies over the past 5 years have documented differences across populations in the number of heterozygous deleterious genotypes, numbers of homozygous derived deleterious genotypes, number of deleterious segregating sites and proportion of sites that are potentially deleterious. These differences have been attributed to population history affecting the ability of natural selection to remove deleterious variants from the population. However, recent studies have suggested that the genetic load may not differ across populations, and that the efficacy of natural selection has not differed across human populations. Here I show that these observations are not incompatible with each other and that the apparent differences are due to examining different features of the genetic data and differing definitions of terms.

## INTRODUCTION

Current evidence from studies of experimental evolution and patterns of genetic variation in natural populations suggest that most new amino-acid changing (i.e. nonsynonymous) mutations are evolutionarily deleterious, and result in a decrease in the number of offspring in individuals who carry them [1,2]. Given their ubiquity, understanding patterns of deleterious mutations are important for interpreting genetic variation and understanding the evolutionary process. Additionally, it has been hypothesized that genetic variants increasing risk to common diseases have been evolutionarily deleterious [3]. Thus, knowledge about the evolution of deleterious mutations will be important to inform studies of the genetics of complex traits.

While strongly deleterious mutations are quickly eliminated from the population by negative natural selection, the behavior of weakly deleterious mutations is more variable. Deleterious mutations, where the strength of selection is less than the inverse of the effective population size, are termed “nearly-neutral”, because, although they are deleterious, they behave in the population as though they are almost neutral, and can reach appreciable frequency in the population [4-8]. The behavior of a particular nearly-neutral mutation is dependent on the population size [6,8]. In large populations, selection will be more effective at reducing the frequency of such a variant. In small populations, however, such mutations can drift to higher frequency, resulting in an apparent decrease in the efficacy of selection.

Because many nonsynonymous mutations in humans are weakly deleterious, it is important to examine their behavior in the context of human population history. Studies of genetic variation suggest that non-African populations have experienced one or more population bottlenecks during the migration process out of Africa [9-12]. Further, recent studies using large samples of individuals suggest that European populations have experienced at least a 100-fold expansion within the last 5,000 years [13-16]. African populations, on the other hand, have not experienced a population bottleneck, and may have experienced ancient growth as well as less-severe recent growth [9-11,14].

Thus, given the ubiquity of deleterious mutations, the potential importance of population history on how natural selection operates, combined with the differences in demographic history across different human populations, it is natural to examine how deleterious mutations are distributed across different human populations. Further, it is also important to assess the mechanism by which specific differences in human history may be affecting natural selection. Over the past 5 years, there have been empirical and theoretical studies examining deleterious variation in different human populations. Several of the studies initially appear to reach disparate conclusions regarding the effects of population history on deleterious genetic variation. Here I review these studies and suggest that they are not incompatible with each other. Apparent discrepancies across studies are due to differences in the statistics used to quantify deleterious variation and subtle variations in the definitions of terms.

## DEMOGRAPHY AFFECTS DELETERIOUS VARIATION

The distribution of deleterious variants in different human populations has been a topic of intense research over the past decade. One of the earlier studies examined PCR-based exon sequencing of just over 10,000 genes in 15 African American (AA) individuals and 20 European American (EA) individuals [17]. This study reported several striking differences in patterns of deleterious variants between the two populations. First, AA individuals had, on average, 1.31 times more nonsynonymous heterozygous genotypes per individual than the EAs. This pattern is likely reflective of the fact that AA populations have had historically larger effective population sizes than European populations [18,19]. The trend was reversed, however, for the number of homozygous genotypes for the derived allele (here defined as the non-chimp allele) per individual. Here, for nonsynonymous variants, EA individuals carried 1.35 times more homozygous derived genotypes per individuals than AA individuals, reflecting variants that reached high frequencies during the out-of-African bottleneck in non-African populations.

The Lohmueller et al. study [17] also compared the distribution of the numbers of putatively neutral variants and putatively deleterious variation in the sample of individuals from both populations. As expected, the AA sample carried more synonymous and nonsynonymous variants than the EA sample. However, the key finding in the Lohmueller et al. study [17] was the proportion of nonsynonymous variants in the EA sample was significantly higher than the proportion of nonsynonymous variants in the EA sample, particularly for population-specific variants. This result suggests that, given the reduction in levels of neutral diversity in the EA sample associated with the Out-of-Africa bottleneck and the levels of nonsynonymous variation in the AA sample, the EA sample contains more nonsynonymous variants than expected. Using forward in time population genetic simulations, Lohmueller et al. [17] showed that this difference in the proportion of nonsynonymous SNPs between the populations is expected. It can be explained by the differences in demographic history experienced by the two populations affecting the ability of negative natural selection to remove deleterious mutations. More recent work has shown that recent population growth is also predicted to increase the proportion of nonsynonymous SNPs [20].

Other recent papers have noted important trends regarding how population history has affected patterns of deleterious mutations. First, several groups have replicated the trends seen by Lohmueller et al. [17]. See Table 1 for a summary of these studies. Another recent paper [21] has estimated the ages of deleterious variants that are currently segregating in populations. The average age of computationally predicted deleterious variants was estimated to be 3,000 years in European Americans and 6,200 years in African Americans [21]. The observation that most deleterious mutations are younger in the European population was likely driven by the influx of new mutations during recent population growth in Europe. Further, Fu et al. [21] found that, for essential genes and genes involved in Mendelian disorders, the proportion of deleterious SNPs decreased monotonically as a function of age in the African population. However, in the European population, the proportion of deleterious SNPs did not monotonically decrease as a function of age. Rather, a higher than expected proportion of the variants inferred to have arisen 50,000 to 100,000 years ago are predicted to be deleterious. Simulations suggest that this pattern can be caused by the population bottleneck increasing the probability that deleterious variants persist in the population [21].

**Table 1.**
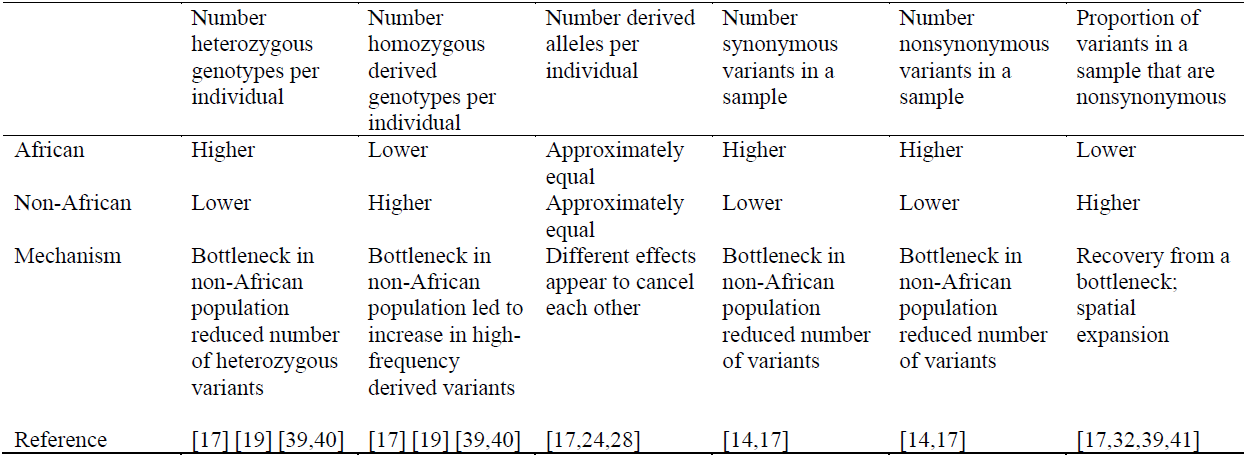
Relative trends of patterns of deleterious variants in African and non-African populations

In addition to genome-wide comparisons across populations, another recent study also examined patterns of deleterious mutations in runs of homozygosity (ROHs) in six African and Native American human populations [22]. ROHs were found to contain a higher proportion of predicted damaging variants than regions of the genome outside of ROHs. Further, the enrichment of damaging variants was stronger for longer runs of homozygosity, suggesting that recent inbreeding can enhance the presence of deleterious genetic variants in the homozygous state, more than would be predicted based upon patterns of neutral variants. Finally, they noted that Native American populations have a higher proportion of predicted damaging variants genome-wide relative to African populations [22]. Taken together, these findings suggest that recent inbreeding has increased the persistence of deleterious mutations in the homozygous state, and there has not been sufficient time for negative selection to eliminate these variants.

Patterns of deleterious variants also appear to differ across populations that have recently split from each other [23]. For example, French –Canadian Quebec was founded by a small number (8,500) of French settlers less than 20 generations ago [23]. This population then expanded more than 700% to its present-day size. An exome sequencing study showed that the French-Canadian population has lower genetic diversity than the French population, supporting the founder-effect suggested by historical records [23]. However, the French Canadian population has a higher proportion of nonsynonymous variants than the French population, particularly at low frequency variants. Additionally, for a given allele frequency, the variants in the French-Canadian population tended to be more disruptive of sites constrained over long evolutionary timescales (i.e. they had higher GERP scores) than the variants in the French population. This finding suggests that nonsynonymous variants segregating in the French-Canadian population are more deleterious than those from the French population [23]. Overall, these results indicate that the very recent differences in demographic history have reshaped patterns of deleterious variants between the two populations.

## A LOAD OF MUTATIONS

Recently Simons et al. [24] reported that differences in human demographic history have “probably had little impact on the average burden of deleterious mutations.” Initially, such a provocative conclusion would seem to be at odds with the studies discussed above documenting differences in patterns of deleterious variants across populations. However, upon more careful examination, these studies are not incompatible with each other. The key insight is that the conclusions from Simons et al. [24] apply to a very particular feature of genetic variation—the genetic load. The genetic load is the reduction in the mean fitness of the population due to the segregation of deleterious variants relative to the fitness of a population having the maximal fitness [25,26]. If the population with the maximum possible fitness has a fitness of 1, then the genetic load (*L*) is 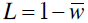 Back where 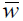 is the mean fitness of the population. Assuming that a deleterious mutation in the homozygous state reduces an individual’s fitness by *s* and reduces an individual’s fitness in the heterozygous case by *hs*, 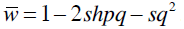.

Simons et al. [24] performed simulations under different models of human population history and distributions of selective effects for deleterious mutations and found that the overall genetic load is predicted to be similar in populations that have experienced a bottleneck and in populations that have experienced recent population growth. Another recent study, using a different simulation strategy, also found that the average genetic load may not differ across populations with different population size changes [20]. These simulations predict that recent population growth results in an influx of new deleterious mutations into the population. These variants tend to be more strongly deleterious, because they are new and selection has not yet had sufficient time to eliminate them. However, these variants also tend to be rare. On the other hand, recent population growth increases the per generation efficacy of selection to reduce the frequencies of pre-existing deleterious variants [27]. These two effects tend to cancel to give the same genetic load in expanded and non-expanded populations.

While the average load may not have been affected by demography, the manner in which a particular population arrives at that load, however, is very dependent on demography (Fig. 1). Specifically, more of the load in the expanded population is accounted for by mutations that tend to be at lower frequency. The load in a population that has not expanded comes from a smaller number of deleterious variants that are each more common in the population. In sum, while the overall genetic load may not have been affected by differences in demography across populations, patterns of deleterious variation are, in fact, affected by demographic history differently than neutral variants.

**Figure 1.**
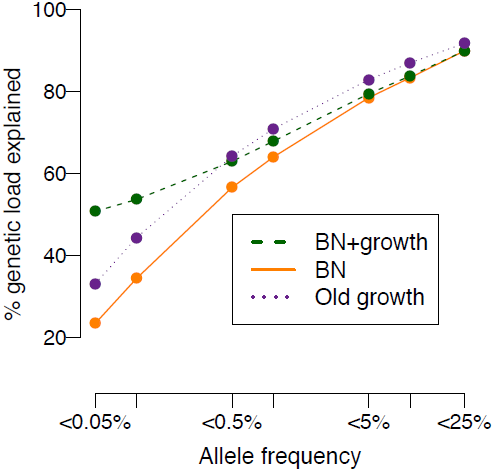
Predicted percent of the genetic load in each population that is accounted for by variants with an allele frequency less than the value specified on the x-axis. BN + growth denotes a model that includes an ancient bottleneck, followed by 100-fold population growth within the last 2,000 years. BN refers to a model with a population bottleneck, without recent growth. Old growth refers to a model with an ancient 2-fold expansion. For a further descriptions of the models, see [20]. Data are taken from the simulations in [20]. Note that low-frequency variants <0.5% account for a greater fraction of the genetic load in a recently expanded population.

Simons et al. [24] also show that the mean derived allele frequency of putatively deleterious variants is similar between African and European populations, and that the number of derived alleles carried by individuals is similar between African and European populations. Such a result is not surprising in light of previous studies using smaller sample sizes [17,28]. However, while the average number of alleles per individual may be similar across populations, how those alleles are distributed into heterozygous and homozygous genotypes has been shown to differ across populations [17]. The extent to which this affects the genetic load depends upon whether mutations are additive (see below).

While genetic load seems like an intuitive measure of the burden of deleterious mutations, this is not necessarily the case. First, the genetic load is a function of the parameters in a population genetic model, and is not something that can be directly empirically measured from genetic variation data. Thus, it is intrinsically difficult to work with. Second, the computation of the genetic load makes particular assumptions about how deleterious mutations interact with each other [26,29]. Typically, mutations are assumed to interact additively or multiplicity, but the biological realism of such a model is not clear, particularly if epistasis is common [26,30]. Third, the computation of the genetic load rests on assumptions about the distribution of fitness effects and dominance coefficients. Just because two populations have similar mean derived allele frequencies does not mean that the load is necessarily similar. The load could differ if the distribution of fitness effects for segregating variants in one population was not the same as that in the other (as suggested by data and simulations [2,17,20]). Further, if mutations are partially recessive, then even if two populations have the same mean derived allele frequency, the load would be predicted to be higher in the population that had more homozygous derived genotypes per individual (because recessive mutations would have a fitness consequence only in the homozygous state). Finally, the relevance of comparing a population with deleterious mutations to a theoretical population free of such mutations has recently been called into question [31].

Even more importantly, the prediction that the genetic load has not been altered by recent human history may not apply to all demographic models. Specifically, in spatial expansion models, the genetic load is predicted to be higher on the expanding wave front than in the ancestral population [32]. This is due to repeated founder effects where deleterious alleles can drift to higher frequency, combined with repeated episodes of population growth increasing the input of deleterious mutations into the populations. Further work with more populations is required to determine which set of models best explains patterns of deleterious variants in multiple human populations.

## EFFICACY OF NATURAL SELECTION

A recent paper has suggested that there is no difference in the efficacy of natural selection across different human populations [33]. Such a conclusion may appear to be at odds with the finding of a higher proportion of nonsynonymous variants in non-African populations compared to what would be predicted from African populations.

To reach their conclusion, Do et al. [33] develop a new statistic that examines the accumulation of deleterious derived mutations on a single chromosome sampled from each of two populations. Using this test, they did not find an excess of deleterious variants on chromosomes taken from non-African populations relative to African populations. However, as shown by their simulations, this test does not have sufficient power to detect differences in the interaction of selection and demography when using distributions of selective effects and demographic models realistic for human populations [33]. Intuitively, this behavior is expected, as this test is based on comparing genetic variation from two chromosomes. Genetic variants detected when comparing two chromosomes are likely to be more common in the population [34], and are more likely to be neutral, rather than deleterious. Further, this test examines the accumulation of intermediate frequency mutations since the two populations split from each other. Because the Out-of-Africa bottleneck occurred relatively recently [9-12], there has not been sufficient time for new mutations to occur and reach intermediate frequency. Thus, the failure to detect a significant difference in the accumulation of deleterious variants using this test does not call into question any of the previous studies (e.g. those in Table 1) that have detected differences in the interaction of demography and selection using other, more powerful, statistics.

Do et al. also examine the behavior of the proportion of nonsynonymous SNPs segregating in a sample [33]. Their simulations show that the proportion of nonsynonymous SNPs is expected to increase after the recovery from a population bottleneck, as previously shown [17] (Fig. 2). However, they perform additional simulations to argue that the increase in the proportion of nonsynonymous SNPs was driven by “neutral forces” rather than the “reduced efficacy of selection” [33]. Because the current population size after the recovery from the bottleneck is actually larger than that during or before the bottleneck, selection is actually more effective at reducing the frequency of a given deleterious variant. In other words, because the effective population size is larger after the population recovered from the bottleneck, it is less likely that a given deleterious mutation would drift to higher frequency. This forms the basis of Do et al.’s conclusion that “the efficacy of selection” has not been reduced in non-African populations [33]. However, this narrow definition of the “efficacy of selection” only considers the efficacy of selection on a given variant, and does not account for the fact that the recent population expansion introduces many new deleterious mutations into the population. This increase in the number of deleterious mutations that enter the population as a result of the population expansion is considered a “neutral force” by Do et al. [33].

**Figure 2.**
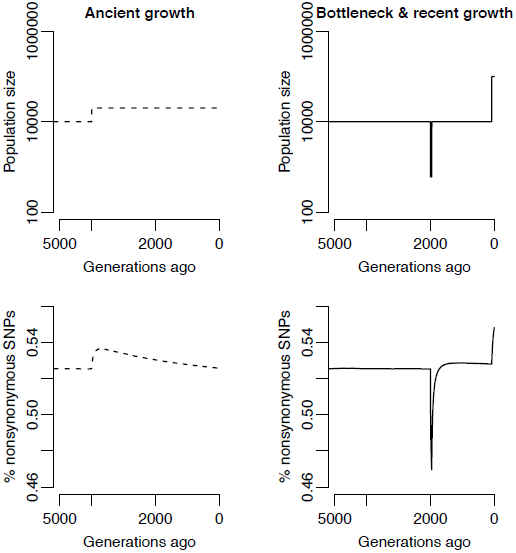
Changes in population size (top) and the proportion of nonsynonymous variants (bottom) for a population with ancient growth (dashed lines; plausible model for African population history) and a population with recent growth (solid lines; plausible model for European population history). Note that the per-generation strength of selection (2*Ns*) is proportional to the population size. 2*Ns* is predicted to be higher in the present-day European population than in the African population, suggesting that, in a given generation, selection acting on a given variant is stronger in the European population. However, the population-scaled deleterious mutation rate is also proportional to the population size. As such, more deleterious mutations recently entered the European population than the African population. Selection has had fewer generations to eliminate them, accounting for the higher proportion of nonsynonymous variants in the expanded population. Figure re-drawn from the simulations in Figures 1 and 2 of [20].

The distinction between “reduced efficacy of selection” and increased importance of “neutral forces” suggested by Do et al. [33] is an unnecessary dichotomy and does not help to clarify the role of population history at shaping deleterious mutations in different populations. Historically, reduced efficacy of selection was taken to mean, by definition, that deleterious mutations behave more like neutral mutations [5,6,8]. Their fate in the population would be determined predominately by mutation and drift, rather than selection. Thus, Do et al.’s [33] conclusion suggests that neutral forces play a greater role in shaping deleterious genetic variation in non-African populations than in African populations. This conclusion would then actually argue, by definition, that selection has been less effective at removing weakly deleterious mutations from non-African populations, due to the increased importance of neutral forces over selective forces.

## CONCLUSIONS AND FUTURE DIRECTIONS

The distribution of deleterious variants in different human populations has been an important topic of research over the past few years. Some summaries of genetic variation, like the proportion of nonsynonymous SNPs, the number of heterozygous genotypes and number of homozygous derived genotypes per individuals show striking differences across populations. Differences in population history, affecting how natural selection removes deleterious mutations from the populations can account for these patterns. Other summaries of genetic variation, like the average derived allele frequency or the number of derived alleles per individual show less striking differences between African and non-African populations.

Despite these clear patterns, much remains to be done. Future work should include examining empirical patterns of deleterious mutations in other human populations that have differing populations histories, such as different amounts of recent population growth. Studies with large samples of individuals will be particularly helpful as they will be informative regarding how deleterious mutations have behaved during recent times [14,15,21,35,36]. Such analyses should better distinguish between whether recent populations growth or a small population size contributes the most to differences in deleterious variants across populations. As the ability to collect and analyze genetic variation data increases, researchers are starting to analyze patterns of deleterious mutations in non-human species with wider range of demographic histories and life-history traits. For example, recent studies found a higher proportion of nonsynonymous SNPs in the self-fertilizing *Capsella rubella* compared to the outcrossing *Capsella grandiflora*, suggesting the recent transition to self-fertilization decreased the efficacy of purifying natural selection [37,38]. Comprehensive analyses of this type in additional species will reveal which aspects of demographic history are most important at affecting the ability of natural selection to remove deleterious mutations from populations.

Additional theoretical and model-based work is also clearly needed for studying deleterious mutations. First, the analyses of Lohmueller et al. [17] and Do et al. [33] point to the importance of non-equilibrium demographic models at affecting deleterious variation. Additional theoretical work on the behavior of deleterious mutations in non-equilibrium populations would greatly contribute to interpreting these patterns. Further, it will be important to examine how spatially explicit models which include deleterious mutations, like those proposed in Peischl et al. [32], fit patterns of genetic variation data across multiple human populations. If such models fit multiple aspects of the data from different populations, it would suggest that the genetic load could actually differ across different populations. These and other future advances in theoretical and empirical work promise to yield even more insights regarding the persistence and distribution of deleterious variants across populations.

## ACKNOWLDEGEMNETS

I thank Joshua Akey, Clare Marsden, Diego Ortega Del Vecchyo, David Reich, and Shamil Sunyaev for helpful discussions on these topics and/or comments on the manuscript.

